# REV7 associates with ATRIP and inhibits ATR kinase activity

**DOI:** 10.1101/2025.01.17.633588

**Authors:** Megan Biller, Sara Kabir, Sarah Nipper, Sydney Allen, Yara Kayali, Skyler Kuncik, Hiroyuki Sasanuma, Pei Zhou, Cyrus Vaziri, Junya Tomida

**Affiliations:** Department of Biological Sciences, University of North Carolina at Charlotte, Charlotte, NC 28223, USA; Department of Genome Medicine, Tokyo Metropolitan Institute of Medical Science, Tokyo 156-8506, Japan; Department of Biochemistry, Duke University School of Medicine, Durham, NC 27710 USA; Department of Pathology and Laboratory Medicine, University of North Carolina, Chapel Hill, NC 27599, USA; Lineberger Comprehensive Cancer Center, University of North Carolina, Chapel Hill, NC 27599, USA

**Keywords:** REV7, CHK1, ATR, DNA repair, ATRIP

## Abstract

Integration of DNA replication with DNA repair, cell cycle progression, and other biological processes is crucial for preserving genome stability and fundamentally important for all life. Ataxia-telangiectasia mutated and RAD3-related (ATR) and its partner ATR-interacting protein (ATRIP) function as a critical proximal sensor and transducer of the DNA Damage Response (DDR). Several ATR substrates, including p53 and CHK1, are crucial for coordination of cell cycle phase transitions, transcription, and DNA repair when cells sustain DNA damage. While much is known about ATR activation mechanisms, it is less clear how ATR signaling is negatively regulated in cells. Here, we identify the DNA repair protein REV7 as a novel direct binding partner of ATRIP. We define a REV7-interaction motif in ATRIP, which when mutated abrogates the REV7-ATRIP interaction *in vitro* and in intact cells. Using *in vitro* kinase assays, we show that REV7 inhibits ATR-mediated phosphorylation of its substrates, including p53. Disruption of the REV7-ATRIP interaction also enhances phosphorylation of CHK1 at Ser317 (a known ATR target site) in intact cells. Taken together our results establish REV7 as a critical negative regulator of ATR signaling. REV7 has pleiotropic roles in multiple DDR pathways including Trans-Lesion Synthesis, DNA Double-Strand Break resection, and p53 stability and may play a central role in the integration of multiple genome maintenance pathways.

## INTRODUCTION

Integration of DNA replication with DNA repair, cell cycle progression and other biological processes is crucial for preserving genome stability and fundamentally important for all life (van Gent *et al*, 2001; Wood *et al*, 2005). Failure to integrate these processes leads to a wide variety of developmental defects and diverse diseases ranging from dwarfism, microcephaly, progressive bone marrow failure, malignancy, and intellectual disability (Bluteau *et al*, 2016; Cimprich & Cortez, 2008; Hira *et al*, 2015; Matsuura *et al*, 1998; Ogi *et al*, 2012; Xu *et al*, 2021; Zhang *et al*, 2011).

The DNA Damage Response (DDR) consists of multiple pathways that regulate DNA repair, DNA replication, and cell cycle progression. Key mediators of the DDR include members of the phosphoinositide 3-kinase related kinase (PIKK) family such as ataxia-telangiectasia mutated (ATM), Ataxia-telangiectasia mutated and RAD3-related (ATR), and DNA dependent protein kinase catalytic subunit (DNA-PKcs). These proteins are serine/threonine kinases that recognize S/TQ motifs on their substrates (O’Neill *et al*, 2000). Members of the PIKK family contain three conserved domains: FRAP-ATM-TRRAP (FAT), FAT carboxy-terminal (FATC), and PIKK regulatory domain (PRD) (Lempiainen & Halazonetis, 2009). FAT domains are thought to be important in cell transport. The small FATC domain is located on the C-terminus and is necessary for kinase activity. The PRD is the PIKK regulatory domain positioned between the FAT and FATC domains, which enhances kinase activity (Lovejoy & Cortez, 2009). All three of these domains are thought to be important for protein-protein interactions (Lempiainen & Halazonetis, 2009).

ATR is a critical proximal sensor and transducer of the DDR (Blackford & Jackson, 2017; Cimprich & Cortez, 2008). While ATM and DNA-PKcs primarily respond to DNA Double-Strand Breaks (DSBs), ATR can be activated by a wider range of genotoxic stress, including DSBs, single-stranded breaks (SSBs), and replication stress (Cimprich & Cortez, 2008). The versatility of ATR and its critical role in maintaining genome integrity during DNA replication makes it indispensable for cell viability, as demonstrated by the embryonic lethality observed in *Atr* knockout mice (Brown & Baltimore, 2000).

ATR activation is a multi-step process that consists of ATR recruitment to sites of damage and localization of one or more activators. The key DNA structure required for ATR activation is single-stranded DNA (ssDNA) coated by replication protein A (RPA) (Zou & Elledge, 2003). ssDNA is exposed after many types of DNA damage. DNA lesions encountered during replication can stall polymerases at the replication fork, resulting in stretches of exposed ssDNA. In addition, DSB end resection can result in ssDNA overhangs. To protect ssDNA from nuclease degradation, it is quickly bound by RPA (Fanning *et al*, 2006). RPA-ssDNA results in the localization of ATR through its obligate subunit ATRIP, which directly binds to RPA and anchors the ATR-ATRIP complex (Zou & Elledge, 2003).

Independently of ATR localization, full activation requires the presence of an activator. Three ATR activators are currently known: TOPBP1, ETAA1, and NBS1. TOPBP1, the main activator of ATR, is directly recruited to sites of damage and requires additional factors, including the RAD17-replication factor C (RFC) complex and the RAD9-RAD1-HUS1 (9-1-1) complex. Localization of TOPBP1 recruits these complexes, and RAD17-RFC loads the 9-1-1 complex onto ss-dsDNA junctions (Yan & Michael, 2009). TOPBP1 directly binds RAD9, ATR, and ATRIP, activating ATR (Ohashi *et al*, 2014). ETAA1 acts independently of TOPBP1 and is recruited to sites of DNA damage by RPA-ssDNA, where it binds RPA through two domains. It then directly binds and activates ATR (Feng *et al*, 2016; Haahr *et al*, 2016). NBS1 has also been shown to directly bind and activate ATR. However, unlike TOPBP1 and ETAA1, NBS1 does not contain an ATR activation domain (AAD), and the mechanisms behind its recruitment are poorly understood (Kobayashi *et al*, 2013).

Once activated, ATR initiates an extensive signaling cascade, with proteomic analyses identifying over a hundred ATR substrates (Matsuoka *et al*, 2007). However, at the core of ATR signaling lie two key substrates: CHK1 and p53. Both CHK1 and p53 play significant roles in DNA repair and cell cycle progression. In response to DNA replication fork stalling, CHK1 signaling can inhibit new origin firing (the S-phase checkpoint) and slows down DNA replication, thereby integrating DNA repair with cell cycle progression (Lukas *et al*, 2004). CHK1 also mediates the G2 checkpoint which ensures complete duplication of the genome before entering mitosis. p53, often referred to as the “guardian of the genome,” plays multiple roles in checkpoint signaling and DNA repair (Lane, 1992). p53 is responsible for initiating the G1 checkpoint, which is critical to preventing the replication of damaged DNA (Helton & Chen, 2007). The genome maintenance roles of CHK1 and p53 are critically ATR-dependent. ATR-mediated phosphorylation of CHK1 on serines 317 and 345 leads to CHK1 activation and triggers the S-phase and G2 checkpoints. ATR-dependent phosphorylation of p53 at serine 15 disrupts p53-MDM2 interactions, thereby stabilizing p53 and promoting its tumor suppressive functions (Tibbetts *et al*, 1999; Zhao & Piwnica-Worms, 2001).

Attenuation of checkpoint signaling is critical for resumption of normal cell cycle progression after DNA damage is resolved. While mechanisms of ATR activation are well described, less is known regarding the negative regulation of ATR activity. Without a brake system, ATR signaling would likely persist unchecked, leading to permanent cell cycle arrest and death. It was recently reported that although ATR may be active in the absence of DNA damage, ATR-CHK1 signaling is suppressed (Li *et al*, 2021). Although a few studies have reported the mechanism of ATR downregulation (Kratz & de Lange, 2018), it remains unclear how cells prevent CHK1 signaling when ATR is active.

In our investigation of novel functions and pathways involving the DNA repair protein REV7, we identified ATRIP as a new REV7 binding partner *in vivo*. We report a novel function of the REV7-ATRIP interaction. Our findings reveal that REV7 directly interacts with ATRIP, leading to the suppression of ATR kinase activity. Through the use of a knock-in system to analyze ATRIP mutants that are incapable of binding REV7, we show that phosphorylation of CHK1 Ser317 is negatively regulated by the REV7 – ATRIP interaction.

## RESULTS & DISCUSSION

### REV7 directly interacts with ATRIP

In a previous study we performed mass spectrometry analysis of REV7 complexes from mammalian cells and identified FAM35A as a REV7-interacting protein (Tomida *et al*, 2018). In the mass spectrometry dataset, we also identified the ATR and ATRIP as components of the REV7 complex (Figure 1A). In an independent approach to verify that ATRIP indeed associates with REV7, we ectopically expressed C-terminally FLAG–HA-tagged ATRIP (ATRIP-FH) or N-terminally FLAG–HA-tagged-ATRIP (FH-ATRIP) in 293T cells. We then immunoprecipitated the two epitope-tagged ATRIP variants using FLAG antibodies and analyzed those complexes using SDS-PAGE and immunoblotting with anti-REV7 antibodies. As shown in Figure 1B, FH-ATRIP and ATRIP-FH both specifically co-immunoprecipitated with endogenous REV7, showing that ATRIP and REV7 exist in the same protein complex in human cells (Figure 1B).

**Figure 1.**
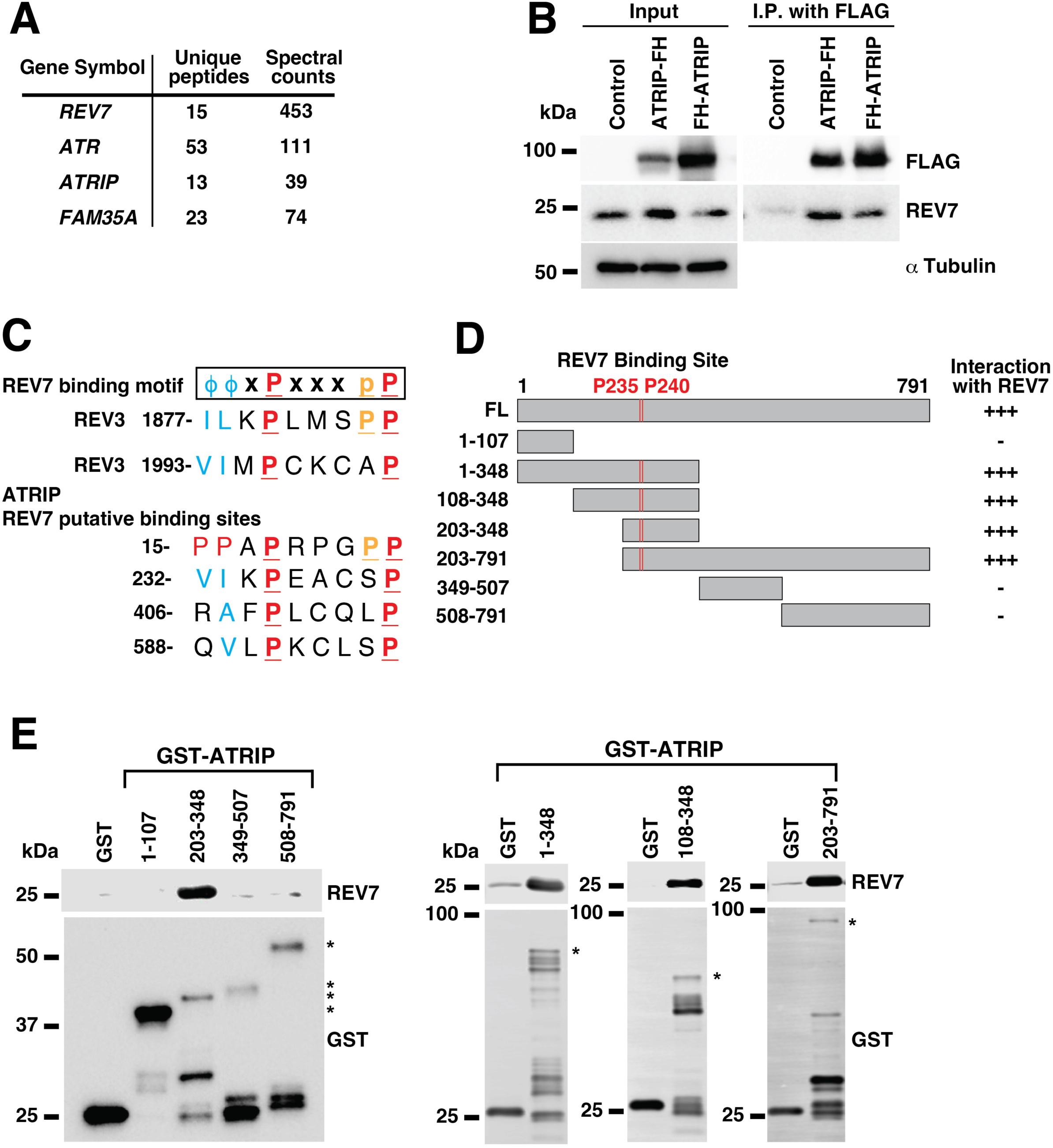
REV7 interacts with the ATR-ATRIP complex. (**A**) REV7 mass spectrometry analysis. The REV7-FH complex was sequentially immunoprecipitated (using FLAG and HA antibody beads) from nuclear extracts of HeLa S3 cell lines stably expressing C-terminally FLAG–HA-tagged REV7 (REV7-FH). A partial list of REV7-FH-associated proteins was identified in the mass spectrometry analysis previously carried out by Tomida et al. The results from the mass spectrometry analysis are shown in the table. A number of unique peptides and Spectral counts (the number of total peptides) were described. (**B**) Co-immunoprecipitation of ATRIP-FH or FH-ATRIP and endogenous REV7 from 293T cells. C-terminally FLAG–HA-tagged ATRIP (ATRIP-FH), (N-terminally FLAG–HA-tagged-ATRIP (FH-ATRIP), or empty vector (control) were transfected into human 293T cells. Thirty hours after transfection, cell lysates were made and used for immunoprecipitation with FLAG antibody beads. After the electrophoretic transfer of proteins, the membrane was immunoblotted with the indicated antibodies. After gel electrophoresis, results for the input and immunoprecipitation (IP) product are shown. (**C**) Sequence alignments of a region of REV3L, showing the REV7 binding sites, and ATRIP with the consensus sequence θ>θ>xPxxxpP at the top. Numbers refer to human ATRIP residues. φ is an aliphatic amino acid, x is any amino acid, P is a highly conserved proline, and p is a less conserved proline. (**D**) *In vitro* association of REV7 with fragments of ATRIP. GST fusion fragments of ATRIP and His-REV7 fusion protein were purified from *E. coli* (see Figure S1). Asterisks (*) indicate the predicted molecular weights of GST fusion fragments of ATRIP. These were used with glutathione beads for GST pulldown experiments. After electrophoresis, samples were immunoblotted with anti-REV7 or anti-GST as indicated. (**E**) Schematic showing locations of the ATRIP fragments GST-ATRIP amino acid residues, 1-107, 1-348, 108-348, 203-348, 203-791, 349-507, and 508-791. The P1235 / P240 position is highlighted in red. See also Supplemental Raw Data Figure 1.

Previous work identified a REV7-binding site ϕϕPxxxxP (where ϕ represents an aliphatic amino acid residue) at residues 1877 – 1887 and 1993 – 2003 of the REV7-binding partner REV3L (Figure 1C). Interestingly, our sequence analyses revealed this consensus REV7-binding sequence motif at residues 232 – 240 in human ATRIP, but not in human ATR (Figure 1C). The REV7-binding sequence in human ATRIP is conserved in *Pan troglodytes* (Chimpanzee), *Macaca mulatta* (monkey), *Canis lupus familiaris* (dog), and Bos Taurus (cattle), but not in Rodentia species (including *Mus musculus* and *Rattus norvegicus* (Figure S2) indicative of a conserved role for ATRIP-REV7 complexes in higher mammals.

To test directly whether the region of ATRIP 204 – 348 can bind to REV7, we purified full-length His-REV7 and several GST fusion fragments of ATRIP in *E. coli* (Figure 1D). We then measured *in vitro* binding of GST-tagged ATRIP fragments with recombinant His-REV7. Our GST pull-down experiments revealed that His-REV7 associated only with GST-ATRIP fragments containing amino acids P235 and P240 (Figure 1E). We conclude that REV7 interacts directly with a 145 amino acid region of ATRIP containing a consensus REV7-interaction motif.

### REV7 interacts with ATRIP at P235 and P240

Next, we asked whether the conserved prolines (P235, P240) in ATRIP function as REV7 binding sites. Therefore, we expressed GST-tagged ATRIP fragments spanning residues 203 – 307 in which these proline residues were conditionally mutated to alanine (P235A, P240A) (Figure 2A). In GST pull-down assays, His-REV7 bound readily to GST-ATRIP 203 – 307 (WT), but not to GST-ATRIP 203 – 307 (P235A, P240A) (Figure 2B). Therefore, P235 and P240 mediate the REV7-ATRIP interaction.

**Figure 2.**
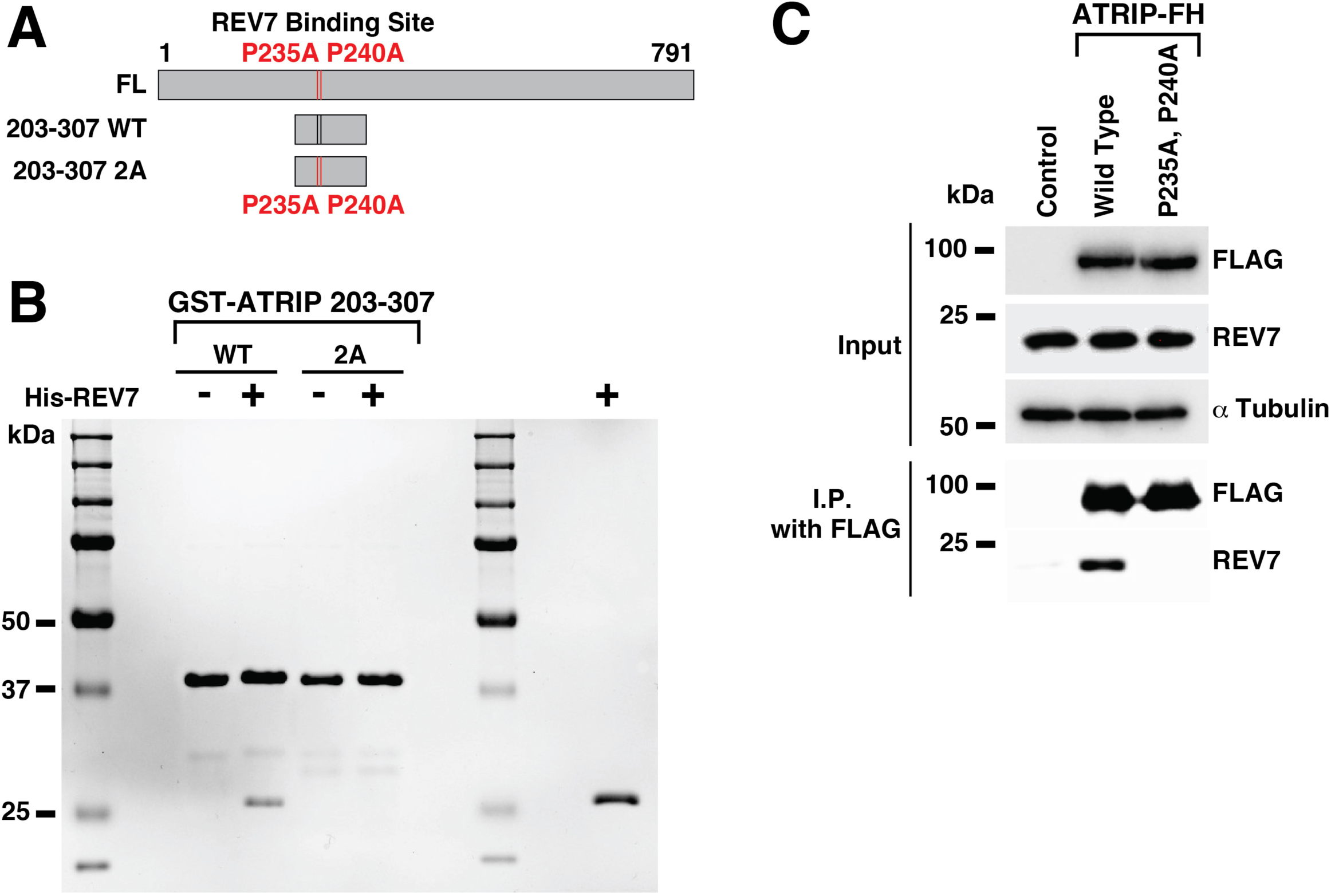
REV7 directly interacts with ATRIP at P235 and P240. (**A**) Schematic drawing showing locations of the ATRIP fragments. The P235/P240 position is highlighted with a vertical bar (red in the online version). (**B**) *In vitro* GST pulldown of purified ATRIP fragments (see Figure S1) containing the indicated amino acid changes. After electrophoresis, samples were stained with Coomassie brilliant blue. (**C**) FLAG-IP assay of 293T cells transfected with ATRIP-FH, ATRIP-FH (P235A, P240A) or control empty vector. Cells were harvested 48 hours after transfection, and immunoprecipitation (IP) was performed using M2 agarose beads. For immunoblotting, we used anti-FLAG, anti-REV7, and anti-αTubulin. See also Supplemental Raw Data Figure 2.

We considered the potential for additional REV7 binding sites in the ATR-ATRIP complex. To test this possibility, 293T cells were transfected with full length ATRIP-FH or with REV7 binding mutants (P235A, P240A). The ATRIP mutant with consensus REV7-binding site substitutions at residues (P235A, P240A) completely failed to bind endogenous REV7 (Figure 2C). Therefore, REV7 interacts directly with ATRIP solely through residues P235 and P240. Interestingly, a previous study established the TOPBP1 binding domain (204 - 308) (Mordes *et al*, 2008), which includes the REV7 binding sites (P235, P240).

### REV7 inhibits ATR kinase activity

Next, we asked whether REV7-ATRIP interactions influence ATR protein kinase activity. Therefore, we performed *in vitro* ATR kinase assays using a recombinant GST-p53 protein (containing residues 2 – 50 which include the ATR target site Ser 15) as a biologically relevant ATR kinase substrate. Since this p53 fragment does not interact with REV7 (Biller *et al*, 2024), the possibility of a REV7-substrate interaction affecting the results is excluded.

As expected, immunoprecipitated FLAG-ATR from 293T cells readily phosphorylated GST-p53 (Figure 3A). Interestingly, FLAG-ATR-dependent phosphorylation of p53 S15 decreased drastically (∼97%) with the addition of purified His-REV7 (Figure 3A), suggesting that REV7-ATRIP interactions inhibit the ability of the ATR complex to phosphorylate its substrates.

**Figure 3.**
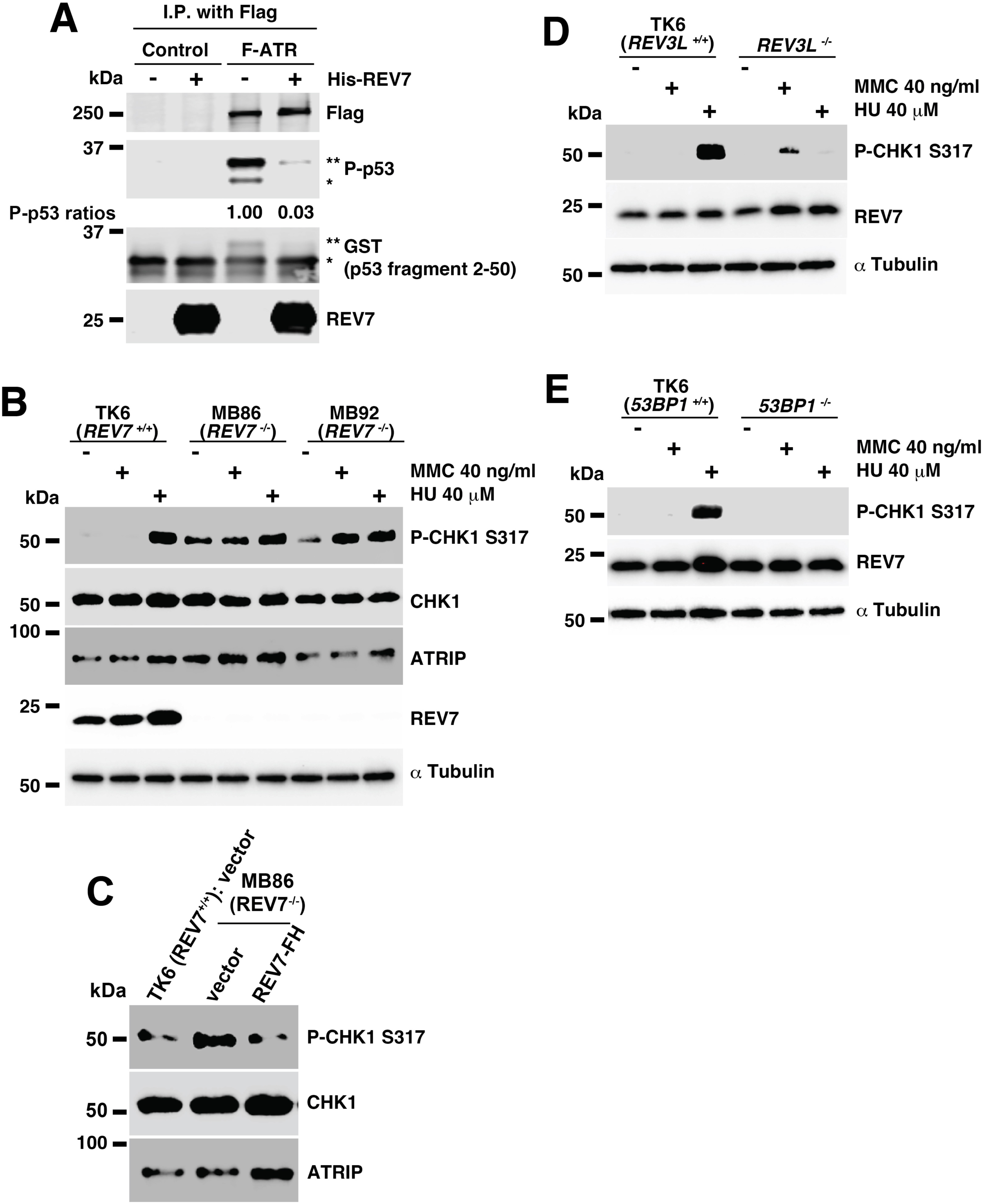
REV7 inhibits ATR activity. (**A**) REV7 disassociated p53 fragment (2 - 50) affects ATR kinase activity *in vitro*. 293T cells were transfected with Flag-tagged ATR (F-ATR) or empty vector control. Cells were harvested 48 hours after transfection, and immunoprecipitation was performed using M2 agarose beads. The beads were washed twice with kinase buffer. 10 μl of buffer or 10 μg of His-REV7 were added, and samples were incubated at 30°C for 15 min. 10 μl of reaction buffer containing substrate (purified GST-p53 fragment) was added followed by a 30 min incubation at 30°C. After electrophoretic transfer of proteins, the membrane was immunoblotted with the indicated antibodies. Results for the immunoprecipitation (IP) product after gel electrophoresis are shown. The shifted P-p53 S15 bands (**) and (*) are the same size as the shifted GST-p53 (2-50) (**) and GST-p53 (2-50) (*) bands, respectively. (**B**) Western blot of DDR activation following a 24-hour 40 ng/ml mitomycin C (MMC), 40 μM hydroxyurea (HU), or mock exposure in TK6 (*REV7*^+/+^), MB86 (*REV7^−/−^*), and MB92 (*REV7*^−/−^) cells. Protein loading was equalized based on cell number. After the electrophoretic transfer of proteins, the Western blot membranes were immunoblotted with the appropriate antibodies. (**C**) The increase in phospho-CHK1 (Ser317) (P-CHK1 S317) protein was rescued when we complemented *REV7*^−/−^ cells with a REV7 expression vector. Protein loading was equalized based on cell number. Expression of CHK1, phospho-CHK1 (Ser317), and ATRIP proteins was determined by Western blot in TK6 (*REV7*^+/+^) with empty vector and MB86 (*REV7^−/−^*) cells with empty vector or REV7-FH. (**D-E**) Western blot of DDR activation following a 24-hour 40 ng/ml mitomycin C (MMC), 40 μM hydroxyurea (HU), or mock exposure in TK6, *REV3L*^−/−^ (**D**) or *53BP1^−/−^* (**E**)TK6 cells. Protein loading was equalized based on cell number. After the electrophoretic transfer of proteins, the Western blot membranes were immunoblotted with the appropriate antibodies. See also Supplemental Raw Data Figure 3.

To test the hypothesis that REV7 is a biologically-relevant negative regulator of ATR signaling, we examined the impact of REV7-loss on the DNA Damage Response (DDR) in intact cells. We used gene editing to generate isogenic WT and *REV7^−/−^* (designated ‘REV7 KO’) TK6 cells. Using these cells, we measured phosphorylation of CHK1 Ser317 following conditional treatment with mitomycin (MMC) or hydroxyurea (HU) as a direct readout of ATR activity. Importantly, REV7-deficiency did not affect expression of ATRIP or CHK1 proteins (Figure 3B). Surprisingly, in *REV7* KO cells, CHK1 Ser317 phosphorylation increased without DNA damage inducing agents (Figure 3B), a signal that was rescued by stable re-expression of REV7-FH (Figure 3C). These findings suggest that REV7 specifically regulates CHK1 Ser317 phosphorylation.

The results of Figure 3 are consistent with our finding that REV7 inhibits ATR, and that REV7 deletion leads to de-repression of ATR kinase activity towards its key substrates CHK1 and p53. We note that REV7 associates with REV3L to form the TLS extender polymerase Polζ, and TLS deficiency has previously been shown to lead to persistent replication stalling and compensatory increases in ATR-CHK signaling (Bi *et al*, 2006; Bi *et al*, 2005). Therefore, we performed additional experiments to determine whether the increased CHK1 317 phosphorylation in *REV7* KO cells could be explained by decreases in Polζ-mediated TLS activity. As an experimental strategy for inhibiting Polζ-mediated TLS independently of REV7-deletion, we tested *REV3L^−/−^* cells. As shown in Fig. 3D, *REV3L* KO cells did not show increases in CHK1 Serine 317 phosphorylation, showing that the increased CHK1 phosphorylation in *REV7* KO cells cannot be attributed to lack of Polζ.

In addition to its role as subunit of Polζ, REV7 is a component of the Shieldin anti-resection complex. Therefore, we also considered the formal possibility that the stimulatory effects of REV7 loss on CHK1 phosphorylation were related to DSB resection. To inhibit DSB anti-resection without depleting REV7 we generated cells lacking 53BP1. As shown in Figure 3E, deletion of *53BP1* had no effect on CHK1 Ser317 phosphorylation without DNA damage induction. Therefore, we conclude that the stimulatory effect of REV7 loss on CHK1 317 phosphorylation cannot be attributed to increased DSB resection.

### Phosphorylation of CHK1 Ser317 is regulated by REV7 – ATRIP binding in intact cells

Next, we sought to verify that the ATR-CHK1 signaling axis is regulated by REV7-ATRIP interactions in intact cells. We co-transfected 293T cells with vectors encoding ATR and WT or REV1-interaction-deficient (P235A, P240A) ATRIP. The ATR/ATRIP-co-expressing cells (and empty vector controls) were conditionally treated with HU to induce S-phase checkpoint signaling. Then we determined the effects of these treatments on levels of CHK1 pSer317 as a measure of ATR activity.

Strikingly, transient co-expression of F-ATR with REV7-interaction-defective ATRIP led to higher levels of CHK1 phosphorylation (approximately two-fold) when compared with cells co-expressing F-ATR with ATRIP (WT) (Figure 4A). We also determined the effect of REV7-depletion on CHK1 Ser317 phosphorylation in cells expressing WT or REV7-interaction-deficient forms of ATRIP. As expected, REV7 depletion led to increases in HU-induced CHK1 Ser317 phosphorylation in cells expressing WT ATRIP (approximately three-fold) (Figure 4B). In contrast, in cells expressing ATRIP (P235A, P240A), CHK1 Ser 317 phosphorylation was relatively insensitive to REV7 depletion (approximately 1.3-fold) (Figure 4B). Taken together, the results of Figure 4 demonstrate that REV7 represses CHK1 Ser317 phosphorylation by ATR, in a manner that depends on REV7-ATRIP interactions in intact cells. Importantly, the repression of ATR signaling by REV7-ATRIP interactions in intact cells, fully recapitulates our biochemical studies with isolated proteins.

**Figure 4.**
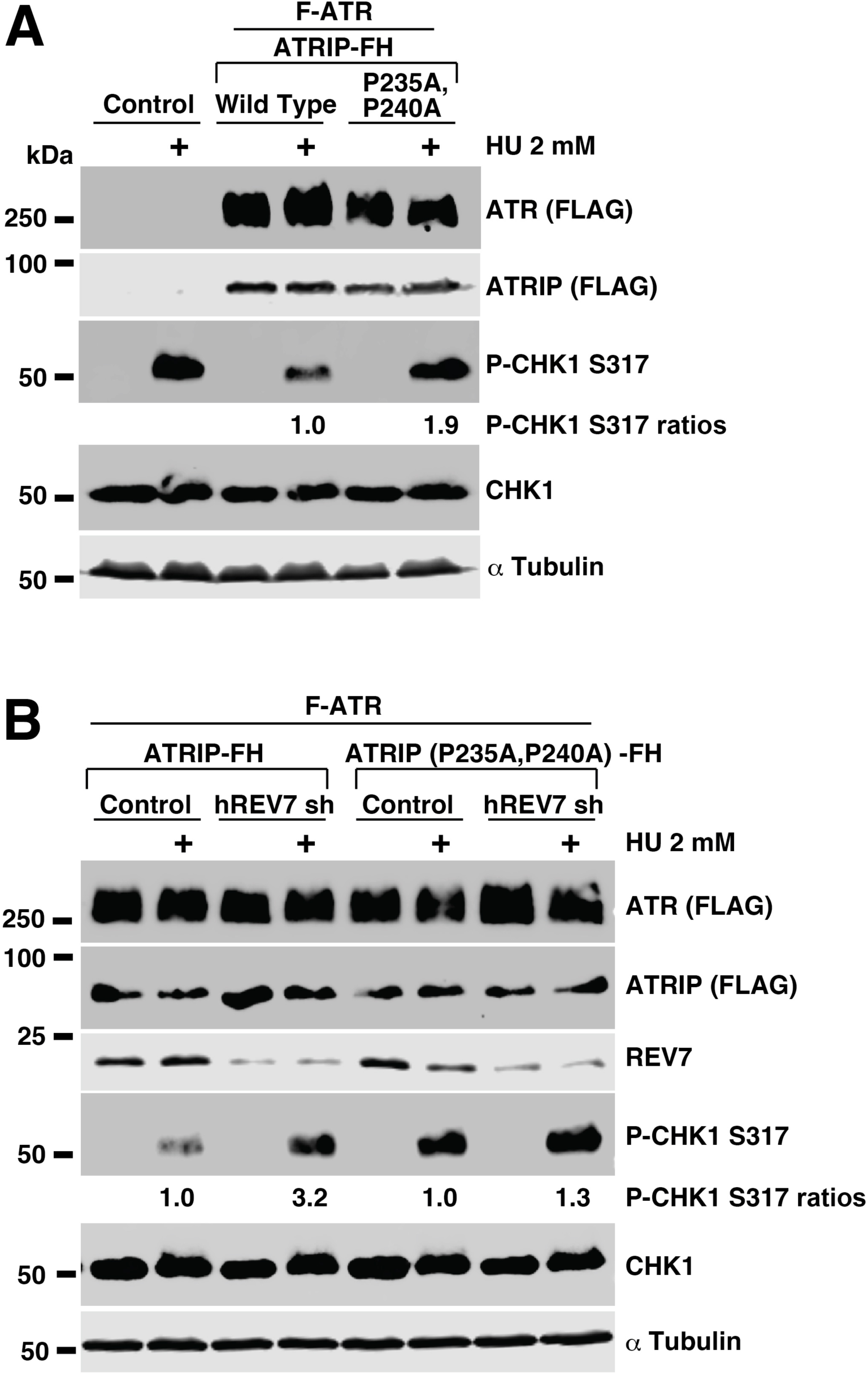
REV7 inhibits Ser317 phosphorylation of CHK1 via ATRIP binding. (**A**) REV7-disassociated ATRIP increases Ser317 phosphorylation of CHK1. FLAG-tagged ATR (F-ATR) was co-transfected into human 293T cells with empty vector (Control), ATRIP-FH, or ATRIP-FH (P235A, P240A). 42 hours after transfection, cells were treated for 6 hours with 2 mM hydroxyurea (HU) or mock. Cells were then harvested and used for Western blot. After the electrophoretic transfer of proteins, the Western blot membranes were immunoblotted with the appropriate antibodies. (**B**) REV7 depletion increases Ser317 phosphorylation of CHK1 in ATR-ATRIP expressing cells but not in cells expressing REV7-disassociated ATRIP. FLAG-tagged ATR (F-ATR) was co-transfected into human 293T cells with ATRIP-FH and shRNA Scramble (Control) or shRNA REV7 (hREV7 sh) and ATRIP-FH (P235A, P240A) and shRNA Scramble (Control) or shRNA REV7 (hREV7 sh). 42 hours after transfection, cells were treated for 6 hours with 2 mM hydroxyurea (HU) or mock. Cells were harvested and used for Western blot. After the electrophoretic transfer of proteins, the Western blot membranes were immunoblotted with the appropriate antibodies. See also Supplemental Raw Data Figure 4.

### CHK1 Ser317 phosphorylation accumulates in cells expressing REV7 binding mutant *ATRIP* without DNA damage induction

To further define the effect of REV7-ATRIP interactions on ATR-CHK1 signaling, and to complement our ATR-ATRIP overexpression studies in 293T cells (Figure 4), we used a knock-in approach to alter endogenous *ATRIP* alleles in diploid TK6 cell lines. Figure S2A describes the gene targeting strategy we used to generate TK6 cells harboring proline (P) to alanine (A) codon substitutions in the endogenous *ATRIP* alleles.

To determine how the REV7-ATRIP interaction affects ATR-CHK1 signaling in TK6 cells, we compared phosphorylation of ATR substrate (CHK1 Ser317) in two independent clones of *ATRIP^P235A, P240A^*TK6 cells and in an isogenic parental cell line. We determined phosphorylation of ATR substrates in all 3 cell lines both basally and after treatment with hydroxyurea (HU).

As expected based on our in vitro experiments and studies with 293T cells, ATRIP^P235A, P240A^ increased CHK1 Ser317 phosphorylation without DNA damage-inducing agents. After HU treatment, phosphorylation of CHK1 Ser317 increased in ATRIP^P235A, P240A^ (Figure 5A). This result suggests that ablation of REV7-ATRIP binding does not affect the TOPBP1-ATRIP interaction, which is necessary for full activation of ATR.

**Figure 5.**
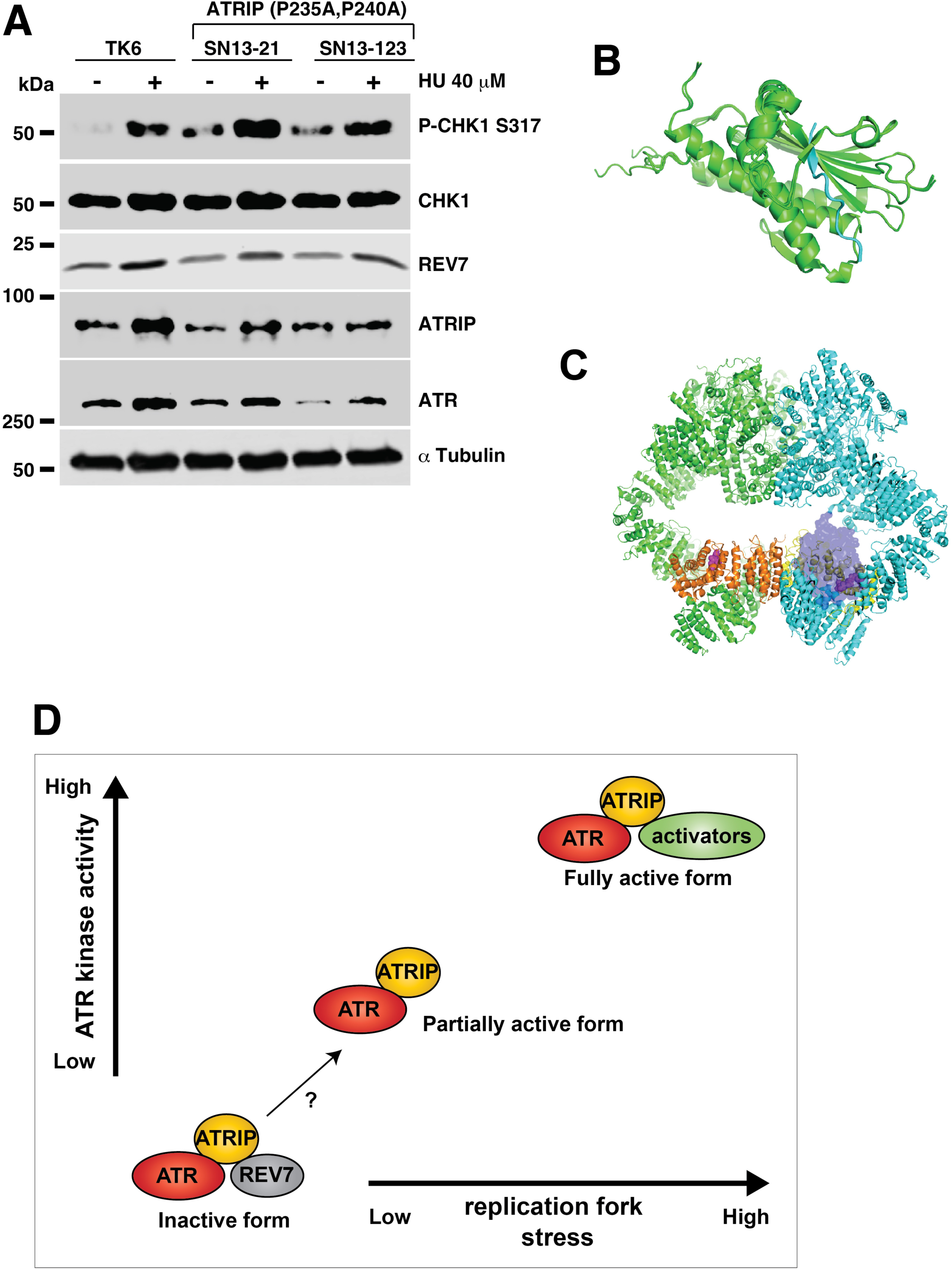
REV7-ATRIP-ATR is the inactive form of the ATR complex. (**A**) Western blot of DDR activation. CHK1 phosphorylation of Serine 317 was examined following a 24-hour exposure to 40 μM hydroxyurea (HU) exposure in TK6, SN13-21 *(ATRIP^P235A, P240A/P235A, P240A^*), and SN13-123 (*ATRIP^P235A, P240A/P235A, P240A^*) cells. Protein loading was equalized based on cell number. After the electrophoretic transfer of proteins, the Western blot membranes were immunoblotted with the appropriate antibodies. (**B**) AlphaFold model of the ATRIP-REV7 complex from ColabFold implementation of the AlphaFold-multimer. Green indicates REV7. Blue indicates REV7 binding site peptide “VIKPEACSPQF” in ATRIP. (**C**) ATR-ATRIP complex with overlay of the REV7 binding site peptide. Green/cyan indicates ATR. Orange/yellow indicates ATRIP. The ATRIP peptide “VIKPEACSPQF” is purple (in space filling). (**D**) Model of functional interactions between REV7 and ATRIP. Bottom left: The REV7-ATRIP-ATR complex is the inactive form. REV7 interacts with ATRIP via P235 and P240 within the ϕϕPxxxxP motif, which is known to bind to the safety belt region of REV7. By binding to ATRIP, REV7 may induce a conformational change in the ATR complex that prevents CHK1 signaling. Functionally, this interaction could be significant, as it may enable REV7 to suppress CHK1 signaling when ATR activation is unnecessary. Middle: Partial complex activation is achieved through the release of REV7. How REV7 dissociates from the ATR-ATRIP complex is unknown. REV7 is capable of dimerizing, and may detach from the complex upon dimerization and/or interaction with REV7 binding proteins. Top right: Full activation of the ATR complex requires association with its activators. The TOPBP1 binding domain (aa 204 - 308) in ATRIP includes the REV7 binding sites (P235, P240). During replication stress TOPBP1 may replace REV7, fully activating the complex. See also Supplemental Raw Data Figure 5.

Finally, we investigated the potential mechanism for the inhibition of ATR-ATRIP kinase activity by REV7. We first generated a structural model of REV7 in complex with an ATRIP peptide (residues 232-242: VIKPEACSPQF) with excellent convergence using the ColabFold (Mirdita *et al*, 2022) implementation of AlphaFold-Multimer (https://www.biorxiv.org/content/10.1101/2021.10.04.463034v2). The structural model was then overlaid with the Cryo-EM structure of the human ATR-ATRIP complex (Rao *et al*, 2018) to evaluate whether the complex provides sufficient space for REV7 binding (Figures 5B and 5C).

Importantly, the ATR-ATRIP complex can phosphorylate CHK1 Ser345 without TOPBP1 (Rao *et al*., 2018). We therefore hypothesized that the Cryo-EM structure of the human ATR-ATRIP complex lacks space for REV7 binding. We found that the REV7 binding peptide resides in a loop region of ATRIP (Figure 5C). Strikingly, this model suggests that the current Cryo-EM structure of the human ATR-ATRIP complex is spatially incompatible with REV7 binding. However, our immunoprecipitation data, GST-pull down results, and ATR kinase assay results all indicate that REV7 interacts with the ATR-ATRIP complex and affects ATR kinase activity.

Altogether, these data suggest that the local structure of ATRIP may undergo a conformational switch to interact with REV7 and then inhibit ATR kinase activity. We conclude that 1) the REV7-ATRIP-ATR complex is the inactive form, 2) the ATRIP-ATR complex is the partially active form, and 3) the complex of ATRIP-ATR with its activators (TOPBP1, ETAA1, and others) is the fully active form (Figure 5D).

Taken together, our results suggest that REV7 is an important negative regulator of the ATR-CHK1-mediated S-phase checkpoint. The best known S-phase function of REV7 involves its association with REV3L to form the TLS extender DNA polymerase, Polζ. Similar to ATR signaling, TLS is initiated by RPA-coated ssDNA. It is generally thought that RPA-ssDNA recruits the RAD18-RAD6 complex to the vicinity of stalled replication forks to promote PCNA mono-ubiquitylation and switching of replicative DNA polymerases with Y-family inserter TLS polymerases. Depending on the nature of the fork-stalling DNA lesion, inserter polymerases may be switched and replaced with Polζ. Jacobs and colleagues have shown that REV7 and REV3 reside at stalled and regressed forks and facilitate both ongoing fork movement and promote protection of HU-regressed forks (Paniagua *et al*, 2022). Many previous studies have demonstrated significant coordination, interplay, and functional redundancy between ATR-CHK1 signaling and TLS. For example historically, many XPV cell lines (which lack functional Polη), were found to be sensitive to UV-induced DNA damage only in the presence of caffeine (Itoh *et al*, 2000; Kannouche *et al*, 2001) which is now known to inhibit ATR. Kannouche and colleagues formally demonstrated a synthetic lethal relationship (i.e. redundancy) between POLH and ATR (Despras *et al*, 2010). TLS-compromised cells lacking RAD18 or inserter Y-family DNA polymerases (such as Polκ or Pol17) fail to recover from replication stalling and exhibit persistent compensatory CHK1 signaling. Thus S-phase checkpoint signaling confers DNA damage tolerance and is induced when TLS fails. Conversely, TLS is important for the timely attenuation of the S-phase checkpoint. In our studies, REV3L-deficiency did not phenocopy the aberrant excessive CHK1 phosphorylation that was induced by REV7-loss. We suggest therefore that REV7 is targeted to ATRIP (to attenuate ATR-CHK1) signaling via a mechanism that does not require Polζ. It is remarkable that the small REV7 protein forms specific complexes with multiple partners to regulate TLS, p53 activity, DSB resection and S-phase checkpoint signaling. Further work is neccesary to clarify the mechanisms by which REV7 is directed to its various partners to facilitate and perhaps integrate diverse genome maintenance processes.

Altogether, our results suggest that the TLS pathway may be an important determinant of ATR substrate choice. Further work is needed to clarify the mechanisms by which the TLS pathway directs ATR substrate selection for activation of the cell cycle checkpoint.

## ACKNOWLEDGEMENTS

We sincerely thank Richard D. Wood (University of Texas MD Anderson Cancer Center) for his early support of this project. We thank Mary Tomida and Luke Tomida for editorial help. **Funding.** This research was supported by National Institutes of Health (NIH) grant 1R15CA263784 (to JT), National Institutes of Health (NIH) grants R01 ES009558, CA215347, and CA229530 (to CV), SPS KAKENHI Grant 19H04267; Takeda bioscience Research grant; Mitsubishi Research grant; Naito Research grant; Astellas research foundation (to HS).

## AUTHOR CONTRIBUTIONS

MB, HS, and JT conceived and designed the experiments. CV and JT designed the research. MB, SK, SN, SA, YK, SK, PZ, and JT performed research. MB, SK, SN, SA, YK, SK, PZ, CV, and JT analyzed data. CV and JT wrote the paper.

---

**Supplemental Figure 1. Purified recombinant GST fusion ATRIP fragments.**

**(A-H**) GST fusion fragments of ATRIP (used to obtain results depicted in Figures 1D and 2B) were expressed in *E. coli*, purified, and stained with Coomassie brilliant blue. Asterisks (*) indicate the predicted molecular weights of GST fusion fragments of ATRIP.

**Supplemental Figure 2. Gene targeting of human ATRIP.**

**(A)** The schema of the ATRIP locus and the ATRIP knock-in vector. The position of mutation sites in Exon 5 is indicated. Red triangles are loxP sites. The first gene-targeting step was to introduce proline (P) to alanine (A) codon substitutions into endogenous *ATRIP* alleles using knock-in vectors, including a puromycin or hygromycin resistance gene cassette (Figure S2A). (**B**) Schematic representation of the gene targeting procedure. We obtained homozygous cell lines harboring P235A and P240A alleles (Figure S2B). Then, cells were transiently co-transfected with Cre recombinase to remove both puromycin or hygromycin resistance gene cassettes (Figure S2B). We obtained two puromycin and hygromycin sensitive clones that exclusively expressed the P235A and P240A mutants (Figure S2B).

**Supplemental Figure 3.** Sequence alignments of a region of ATRIP, showing identified REV7 binding sites, with the consensus sequence SVVIKPEACSPQF at the bottom. Numbers refer to human ATRIP residues. An amino acid alignment of *Homo sapiens* (human) (accession number NP_569055.1), *Pan troglodytes* (Chimpanzee) (XP_001156485.2), *Macaca mulatta* (monkey) (XP_001112538.2), *Canis lupus familiaris* (dog) (XP_005632647.1), and Bos Taurus (cattle) ATRIP (NP_001096800.1) proteins.

## STAR Methods

### RESOURCE AVAILABILITY

#### Lead Contact

Additional information and requests for resources and reagents can be directed to and fulfilled by Lead Contact, Dr. Junya Tomida.

#### Material Availability

This study did not generate unique reagents.

#### Data and Code Availability

All data are presented in the article.

### EXPERIMENTAL MODEL AND SUBJECT DETAILS

#### Cell lines

##### Human cell cultures and transfections

All cells were maintained in a humidified 5 % CO_2_ incubator at 37 °C. HEK293T (ATCC CRL11268) cells were cultured in Dulbecco’s Modified Eagle’s Medium (DMEM) (Life Technologies, Carlsbad, CA) supplemented with 10 % fetal bovine serum and 1 % penicillin/streptomycin (Invitrogen). Human TK6 (WT), *53BP1*^−/−^ (Sasanuma *et al*, 2018), and *REV3L^−/−^* (Saha *et al*, 2018) cell lines were obtained from Dr. Hiroyuki Sasanuma, Tokyo Metropolitan Institute of Medical Science. Human TK6 cells were cultured in RPMI supplemented with 5 % horse serum (Gibco^TM^) and 1 mM sodium pyruvate (Gibco^TM^).

All cell lines were routinely assessed for mycoplasma contamination using the MycoAlert detection kit (Lonza). Cell transfection utilized a polyethylenimine (PEI) method as described previously (Aricescu *et al*, 2006; Biller *et al*., 2024; Lee *et al*, 2014).

### METHOD DETAILS

#### Generation of *ATRIP* knock-in TK6 cells

*ATRIP* guide RNA (gRNA) target sequence (5’-GAAAGCTTGTCTTAAGCTGGAGG -3’) was inserted into the pX330 vector (Addgene, US) for the CRISPR Cas9 system. To generate *ATRIP^P235A, P240A^*for TK6 cells, *ATRIP^P235A, P240A^ -PURO^R^* and *ATRIP^P235A, P240A^ -HYG^R^* targeting vectors, a gift from Dr. Hiroyuki Sasanuma (Kratz *et al*, 2021; Tsuda *et al*, 2017), were produced from PCR-amplified genomic products combined with *PURO^R^* and *HYG^R^* selection marker genes. PCR-amplified genomic products were amplified using the following primers: 5’-GCGAATTGGGTACCGGGCCGCTCCAGCATGGGCAACAGAGTGAGACCCTGTC-3’ and 5’-CTGGGCTCGAGGGGGGGCCAGTGATTCCAGTCCCTGCAGAGATTGTCCCCTC-3’ plus 5’-TGGGAAGCTTGTCGACTTAAATCTATTTCAAGGTGGCTCACTCACATGGCTG-3’ and 5’-CACTAGTAGGCGCGCCTTAACATTTTGGGAGGCTGAAGTGGGTGGATCACG-3’ for the left arm and right arm, respectively. Left and right arms were inserted into *Apa*I and *Afl*II sites of targeting vectors, respectively, using In-Fusion (Takara). All transfections in TK6 were performed as described (Ibrahim *et al*, 2020; Kratz *et al*., 2021; Saha *et al*., 2018; Tsuda *et al*., 2017). Cells were seeded in 96-well plates and allowed to grow. Resultant colonies were screened for *ATRIP^P235A, P240A^ -PURO^R^* and *ATRIP^P235A, P240A^ -HYG^R^* through integration of antibiotic resistance was confirmed. After isolation clones, ATRIP^P235A, P240A^ mutations were confirmed by DNA sequencing. 1 μg pCMV-Cre (Addgene plasmid # 123133) was transfected in 1 × 10^6^ *ATRIP^P235A, P240A^ -PURO^R^* and *ATRIP^P235A, P240A^ -HYG^R^* TK6 cells. Resultant colonies were screened for *ATRIP^P235A, P240A^* through the confirmation of antibiotic sensitivity.

#### DNA damage treatment

TK6 (WT, *REV7*^−/−^, *REV3L*^−/−^, and *53BP1*^−/−^) cell lines were seeded in T25 flasks at a concentration of 2 × 10^5^ cells/mL. Cells were treated with 40 ng/ml mitomycin C (MMC), 40 μM hydroxyurea (HU), or control. 24 hours later, 4 × 10^6^ cells were harvested and resuspended in 200 μl of 1x SDS loading buffer as described previously (Biller *et al*., 2024). Samples were sonicated (30 % amplitude, 4 cycles of 15 seconds with a 30 second pause). 42 hours after transfection, 293T cells were treated with 2 mM hydroxyurea (HU) or mock for 6 hours. Cells were then harvested, resuspended in 200 μl of 1x SDS loading buffer and sonicated (30 % amplitude, 4 cycles of 15 seconds with a 30 second pause).

#### Protein purification, GST pulldown, immunoprecipitation, and immunoblotting

Proteins were purified from *E. coli* as described previously (Biller *et al*., 2024; Takata *et al*, 2006; Takeuchi *et al*, 2004; Tomida *et al*, 2015). His-REV7 and GST purification were performed as described (Biller *et al*., 2024). Immunoprecipitation and immunoblotting were described previously (Takata *et al*, 2013; Tomida *et al*, 2013; Tomida *et al*., 2018). The samples were separated by polyacrylamide gel electrophoresis, transferred to a membrane, and detected with the indicated antibodies and ECL reagents (GE Healthcare) or IRDye 800 (Li-Cor).

#### ATR kinase assays

Kinase assays were performed as described previously (Biller *et al*., 2024).

#### Antibodies

Antibodies purchased along with dilutions used for immunoblotting were as follows: p-p53 S15 (9284), 1:1,000; p-Chk1 S317 (12302), 1:1,000, were obtained from Cell Signaling. MAD2B (612266), 1:1,200, was purchased from BD Biosciences. F3165, monoclonal anti-FLAG 1:10,000; T5168, monoclonal anti-α-Tubulin 1:8,000; A0168 HRP (horseradish peroxidase) conjugated anti-mouse IgG 1:10,000; and A0545 HRP conjugated anti-rabbit IgG 1:10,000 were purchased from Sigma-Aldrich. IRDye 800 CW Goat anti-Mouse IgG Secondary Antibody (926-32210), 1:20,000, and IRDye 800 CW Goat anti-Rabbit IgG Secondary Antibody (926-32211), 1:20,000, were purchased from Li-Cor. Anti-GST (SC-138), 1:400; p53 (sc-126), 1:200; CHK1 (sc-8408), 1:2500; and ATR (sc-515173) 1:200, were purchased from Santa Cruz Biotechnology.

#### DNA constructs

The human His-REV7 (pEtDuet-1-REV7) (Tomida *et al*., 2015), pCDH-EF1α-MCS-Flag-HA-IRES-Puro (Yousefzadeh *et al*, 2014), and the following shRNA vector shREV7#2 (TRCN0000006569) were gifted from Dr. Richard Wood. shScramble shRNA was obtained from Dr. David Sabatini (Addgene plasmid # 1864) (Sarbassov *et al*, 2005). The Flag-ATR expression vectors were a gift from Dr. Karlene A. Cimprich (REF). The human *ATRIP* wild type (WT) full-length cDNA was obtained from Dr. Minoru Takata (Tomida *et al*., 2013). After construction, expression vectors were confirmed by DNA sequencing. p53: GST-p53 2-50 fragment expression vector was described previously (Biller *et al*., 2024). ATRIP: GST-ATRIP fragments were purified as an XhoI–NotI fragment from the ATRIP expression vector and cloned into pGEX6P-1 (GE Healthcare). GST-ATRIP 1-107, 1-348, 108-348, 203-348, 203-791, 203-307, 349-507, 508-791 fragments were PCR amplified from ATRIP-GFP expression vector as a XhoI–NotI fragment with 5′ATRIP (XhoI) primer (5′ - CCGCTCGAGATGGCGGGGACCTCCGCGCCAGGC) and 3′ ATRIP 107 (NotI) primer (5′ - TAAAAGCGGCCGCTCATGGAACAGTTTCTCTGTTTTTC), 5′ATRIP (XhoI) primer (5′ - CCGCTCGAGATGGCGGGGACCTCCGCGCCAGGC) and 3′ ATRIP 348 (NotI) primer (5′ - TAAAAGCGGCCGCTCATGGTGGCTGCAGGGGGGTGCCAG), 5′ATRIP 108 (XhoI) primer (5′ -CCGCTCGAGATTAAAGATAATTTCGAATTAGAG) and 3′ ATRIP 348 (NotI) primer (5′ -TAAAAGCGGCCGCTCATGGTGGCTGCAGGGGGGTGCCAG), 5′ATRIP 203 (XhoI) primer (5′ -CCGCTCGAGAGGACAAAGCTCCAGACCAGTGAAC) and 3′ ATRIP 348 (NotI) primer (5′ -TAAAAGCGGCCGCTCATGGTGGCTGCAGGGGGGTGCCAG), 5′ATRIP 203 (XhoI) primer (5′ -CCGCTCGAGAGGACAAAGCTCCAGACCAGTGAAC) and 3′ ATRIP (NotI) primer (5′ - TAAAAGCGGCCGCTCAGCCACACTCCACCTCGGGGTCTTC), 5′ATRIP 203 (XhoI) primer (5′ -CCGCTCGAGAGGACAAAGCTCCAGACCAGTGAAC) and 3′ ATRIP-307 rev (NotI) primer (5′ - CAGTCAGTCACGATGCGGCCTCAAGTGTTTGATCTCTGTCTCCAGCTG), 5′ATRIP 349 (XhoI) primer (5′ -CCGCTCGAGGGGTTTGGCAGTACCTTGGCTGGAATG) and 3′ ATRIP 507 (NotI) primer (5′ -TAAAAGCGGCCGCTCACCCAGCAGCAGAATCTGCCCC), and 5′ATRIP 508 (XhoI) primer (5′ -CCGCTCGAGGAAGGAAACAGGAGCCTGGTTCAC) and 3′ ATRIP (NotI) primer (5′ -TAAAAGCGGCCGCTCAGCCACACTCCACCTCGGGGTCTTC) to clone into pGEX6P-1 (GE Healthcare). ATRIP (P235A, P240A) mutation was introduced using site-directed mutagenesis (TOYOBO KOD FX). The XhoI–NotI fragments from *WT ATRIP* or *ATRIP* (P235A, P240A) were inserted into pCDH-EF1α-Flag-HA-MCS-IRES-Puro to generate FH-ATRIP, ATRIP-FH and ATRIP (P235A, P240A)-FH (P235A, P240A).

#### Quantification

Histograms from Western blots (Supplemental Raw Data Figure 6) were created using ImageJ software. Signal intensity was calculated by ImageJ software, with the average signal intensity (n=3) used for the calculations in Figure. 3A, Figure 4A and 4B, the signal of phosphorylated CHK1S317 was normalized to CHK1.

#### Mass spectrometric analysis of purified REV7 complexes

Purification of the REV7 complex and analysis by LC-MS/MS has been described (Tomida *et al*., 2018).

#### Molecular Modeling

The sequences of human REV7 (UniProt: Q9UI95) and a peptide of ATRIP (residues 232-242: VIKPEACSPQF) were used as the input for the ColabFold (Mirdita *et al*., 2022) implementation of AlphaFold-Multimer (https://www.biorxiv.org/content/10.1101/2021.10.04.463034v2) to build the AlphaFold2 model (Tunyasuvunakool *et al*, 2021) of the REV7-ATRIP peptide complex, with an excellent convergence for all five models.

