## Supplementary figures and images for "REV7 associates with ATRIP and inhibits ATR kinase activity"

### Supplemental Figures 1-3

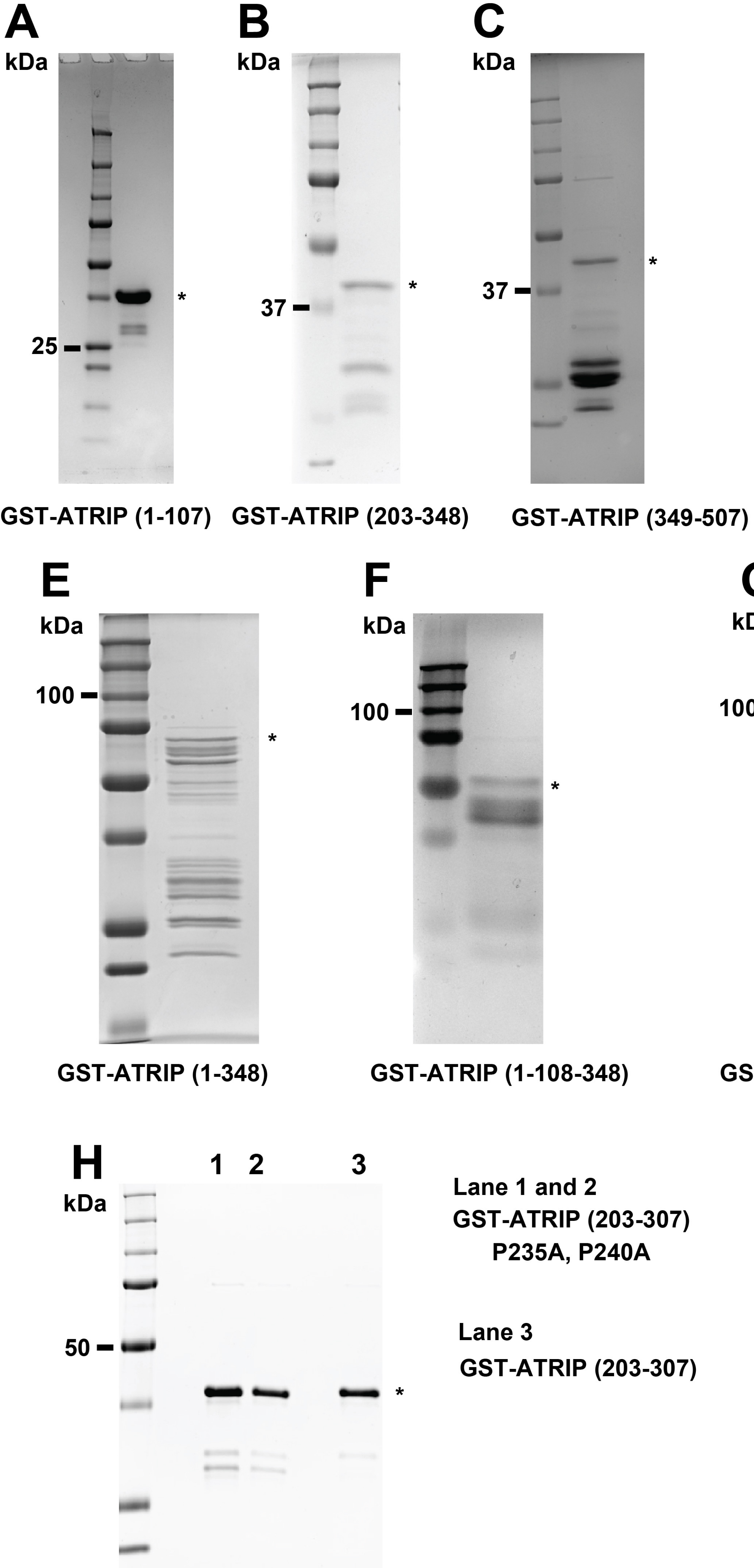


Supplemental Figure 1

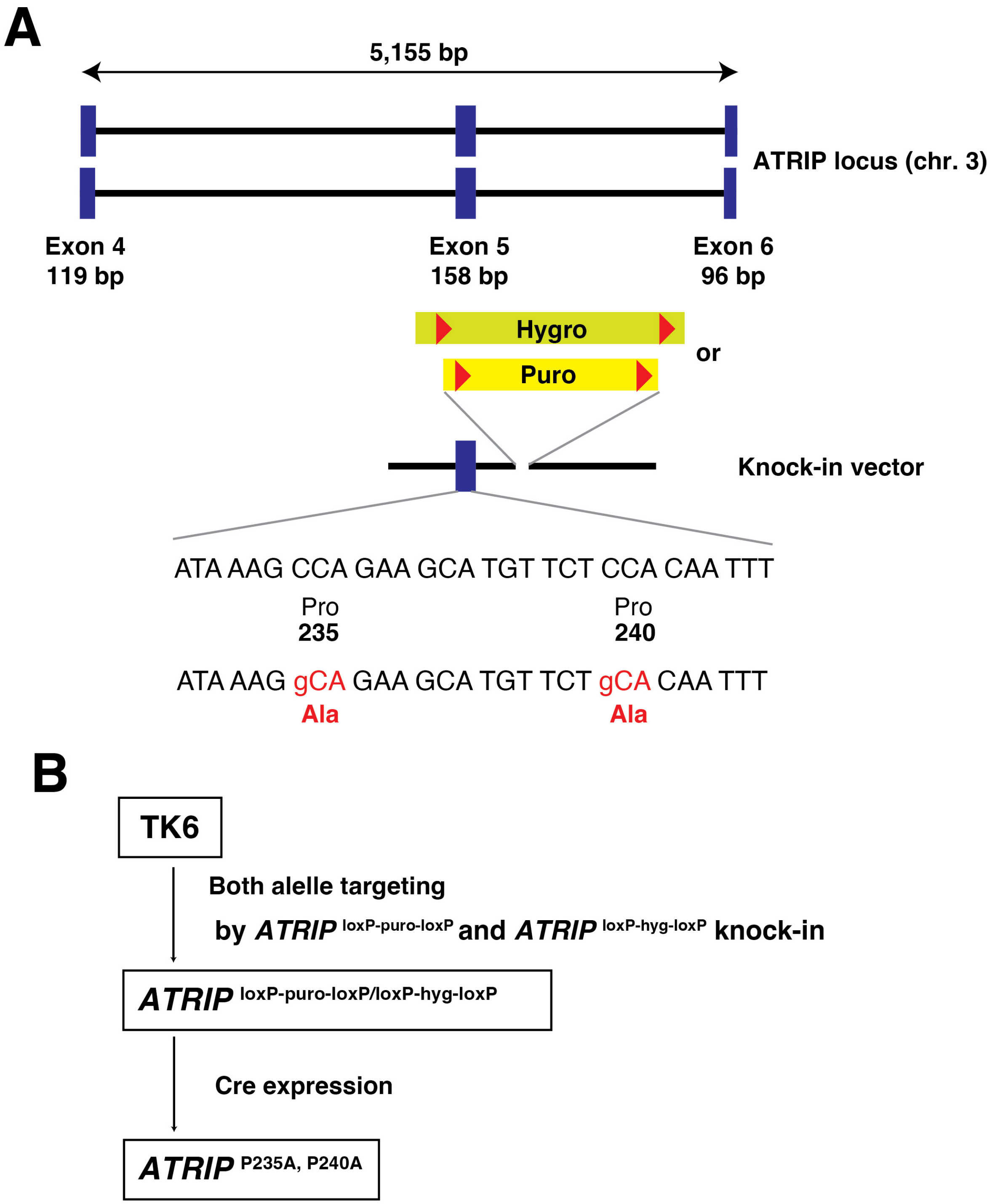


Supplemental Figure 2

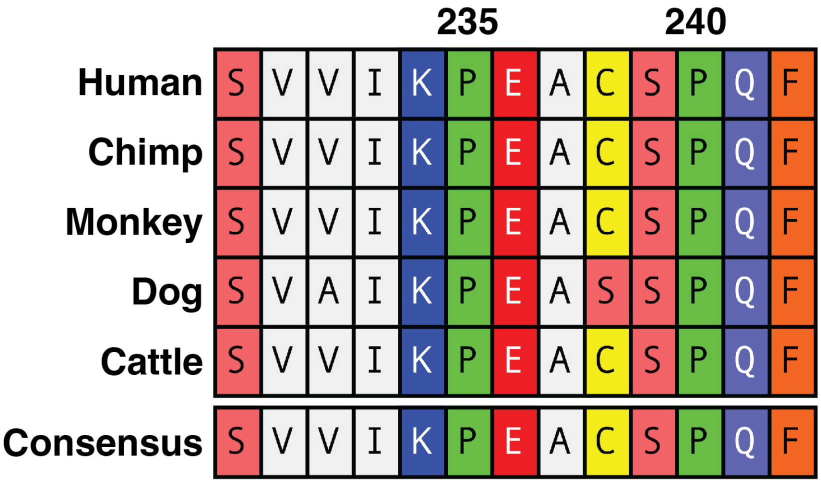


**Supplemental Figure 3**
